# Reduced D-serine levels drive enhanced non-ionotropic NMDA receptor signaling and destabilization of dendritic spines in a mouse model for studying schizophrenia

**DOI:** 10.1101/2021.03.04.434016

**Authors:** Deborah K. Park, Samuel Petshow, Margarita Anisimova, Eden V. Barragan, John A. Gray, Ivar S. Stein, Karen Zito

## Abstract

Schizophrenia is a psychiatric disorder that affects over 20 million people globally. Notably, schizophrenia is associated with decreased density of dendritic spines and decreased levels of D-serine, a co-agonist required for opening of the *N*-methyl-D-aspartate receptor (NMDAR). We hypothesized that lowered D-serine levels associated with schizophrenia would enhance ion flux-independent signaling by the NMDAR, driving destabilization and loss of dendritic spines. We tested our hypothesis using the serine racemase knockout (SRKO) mouse model, which lacks the enzyme for D-serine production. We show that activity-dependent spine growth is impaired in SRKO mice, but can be acutely rescued by exogenous D-serine. Moreover, we find a significant bias of synaptic plasticity toward spine shrinkage in the SRKO mice as compared to wild-type littermates. Notably, we demonstrate that enhanced ion flux-independent signaling through the NMDAR contributes to this bias toward spine destabilization, which is exacerbated by an increase in synaptic NMDARs in hippocampal synapses of SRKO mice. Our results support a model in which lowered D-serine levels associated with schizophrenia enhance ion flux-independent NMDAR signaling and bias toward spine shrinkage and destabilization.

## Introduction

Experience-dependent growth and long-term stabilization of dendritic spines are critical for learning and memory (Hayashi-Takagi et al., 2015). Long-term changes in spine structure and stability are tightly associated with long-term potentiation (LTP) (Hill and Zito, 2013; Matsuzaki et al., 2004) and long-term depression (LTD) (Oh et al., 2013; Zhou et al., 2004) of synaptic strength and are mediated through activation of *N*-methyl-D-aspartate receptors (NMDARs). NMDARs can signal through flux of positive ions, including Ca^2+^, in response to binding of both glutamate and co-agonist, glycine or D-serine. In addition, NMDARs can signal in an ion flux-independent manner upon the binding of agonist or co-agonist alone (Valbuena and Lerma, 2016). Notably, recent studies have established that glutamate binding to NMDARs in the absence of co-agonist drives LTD and spine shrinkage (Nabavi et al., 2013; Stein et al., 2015).

Dysfunctional NMDAR signaling is thought to contribute to the etiology of schizophrenia, a psychiatric disorder that affects up to 1% of the global population and is characterized by a variety of symptoms such as hallucinations and cognitive deficits (Coyle, 2017). These debilitating symptoms may be linked to synaptic and neuroanatomical changes in the brain, such as decreased spine densities (Rosoklija et al., 2000; Sweet et al., 2009). As NMDARs mediate bidirectional structural plasticity and stabilization of spines (Hill and Zito, 2013; Stein et al., 2021), alterations in NMDAR function influence spine densities and impact the ability to learn and form memories (Alvarez et al., 2007; Brigman et al., 2010; Kannangara et al., 2015; Ultanir et al., 2007). Notably, individuals with schizophrenia have decreased levels of D-serine (Bendikov et al., 2007; Hashimoto et al., 2003a) and elevated levels of kynurenic acid (Plitman et al., 2017), an endogenous antagonist of the NMDAR co-agonist site. In addition, the serine racemase enzyme for D-serine production is a risk gene for schizophrenia (Coyle, 2017). Importantly, animal studies that mimic NMDAR hypofunction by reducing ion flux or co-agonist binding show cognitive deficits and decreased spine density (Barnes et al., 2014; Basu et al., 2009; Latysheva and Raevskii, 2003; Schobel et al., 2013; Wu et al., 2016).

We hypothesized that the decreased D-serine level associated with schizophrenia promotes ion flux-independent NMDAR signaling (Nabavi et al., 2013; Stein et al., 2015), creating a bias for spine shrinkage that ultimately leads to decreased spine density. To test our hypothesis, we used the serine racemase knockout (SRKO) (Basu et al., 2009) mouse model, which lacks the enzyme required for D-serine production and exhibits decreased levels of D-serine, decreased spine densities, and cognitive deficits (Balu et al., 2013; Basu et al., 2009). We found that SRKO mice exhibit impaired activity-dependent spine growth, and that activity-dependent spine structural plasticity in SRKO animals is biased toward spine shrinkage and destabilization. Furthermore, we observed increased numbers of synaptic NMDARs and reduced CaMKII activation at synapses of SRKO animals. Finally, we demonstrate that non-ionotropic NMDAR signaling is increased at hippocampal synapses of SRKO animals.

## Materials and methods

### Animals

SRKO (Basu et al., 2009) and GFP-M (Feng et al., 2000) mice in a C57BL/6J background were crossed to generate serine racemase knockout and wild-type littermates with GFP expressed in a subset of hippocampal pyramidal neurons. All experimental protocols were approved by the University of California Davis Institutional Animal Care and Use Committee.

### Two-photon imaging and image analysis

Acute hippocampal slices were prepared from P14-21 WT and SRKO littermates of both sexes as described (Stein et al., 2021). GFP-expressing CA1 pyramidal neurons at depths of 10-50 μm were imaged using a custom two-photon microscope (Woods et al., 2011). For each neuron, image stacks (512 × 512 pixels; 0.02 μm per pixel^;^ 1-μm z-steps) were collected from one segment of secondary or tertiary basal dendrite at 5 min intervals at 27-30 °C in recirculating artificial cerebral spinal fluid (ACSF; in mM: 127 NaCl, 25 NaHCO_3_, 1.2 NaH_2_PO_4_, 2.5 KCl, 25 D-glucose, aerated with 95%O_2_/5%CO_2_, ∼310 mOsm, pH 7.2) with 1 μM TTX, 0.1 mM Mg^2+^, and 2 mM Ca^2+^, unless otherwise stated. Cells were pre-incubated for at least 10 min with 10 μM D-serine or for at least 30 min with 10 μM L-689,560 (L-689), 10 μM Bay-K and 50 μM NBQX (all from Tocris), which were included as indicated. Images are maximum projections of three-dimensional image stacks after applying a median filter (3 × 3) to raw image data. Estimated spine volume was measured from background-subtracted green fluorescence using the integrated pixel intensity of a boxed region surrounding the spine head, as described (Woods et al., 2011).

### Glutamate uncaging

High-frequency uncaging (HFU) consisted of 60 pulses (720 nm; 2 ms duration, 7-11 mW at the sample) at 2 Hz delivered in ACSF containing (in mM): 2 Ca^2+^, 0.1 Mg^2+^, 2.5 MNI-glutamate, and 0.001 TTX. The beam was parked at a point 0.5-1 μm from the spine at the position farthest from the dendrite. HFU+ stimulation consisted of 60 pulses (720 nm; 8 ms duration, 6-10 mW at the sample) at 6 Hz, delivered in ACSF containing (in mM): 10 Ca^2+^, 0.1 Mg^2+^, 5 MNI-glutamate, and 0.001 TTX. For experiments using HFU+ stimulation, healthy and stimulus responsive cells were selected as described (Stein et al., 2021).

### Electrophysiology

Modified transverse 300 μm slices of dorsal hippocampus were prepared from P15–P19 mice anesthetized with isoflurane (Bischofberger et al., 2006), and mounted cut side down on a Leica VT1200 vibratome in ice-cold sucrose cutting buffer containing (in mM): 210 sucrose, 25 NaHCO_3_, 2.5 KCl, 1.25 NaH_2_PO_4_, 7 glucose, 7 MgCl_2_, and 0.5 CaCl_2_. Slices were recovered for 1 hour in 32°C ACSF solution containing (in mM): 119 NaCl, 26.2 NaHCO_3_, 11 glucose, 2.5 KCl, 1 NaH2PO4, 2.5 CaCl_2_, and 1.3 MgSO_4_. Slices were perfused in ACSF at RT containing picrotoxin (0.1 mM) and TTX (0.5 μM) and saturated with 95% O_2_ / 5% CO_2_. mEPSCs were recorded from CA1 pyramidal neurons patched with 3–5 MΩ borosilicate pipettes filled with intracellular solution containing (in mM): 135 cesium methanesulfonate, 8 NaCl, 10 HEPES, 0.3 Na-GTP, 4 Mg-ATP, 0.3 EGTA, and 5 QX-314 (290 mOsm, pH 7.3). Cells were discarded if series resistance varied by more than 25%. Recordings were obtained with a MultiClamp 700B amplifier (Molecular Devices), filtered at 2 kHz, digitized at 10 Hz. Miniature synaptic events were analyzed using Mini Analysis software (Synaptosoft) using a threshold amplitude of 5 pA for peak detection. To generate cumulative probability plots for amplitude and inter-event time interval, events from each CA1 pyramidal neuron (>100 per cell) were pooled. The Kolmogorov-Smirnov two-sample test (KS test) was used to compare the distribution of events between WT and SRKO. Statistical comparisons were made using Graphpad Prism 8.0.

### Biochemistry

Hippocampi of P20 mice of either sex were homogenized with 1% deoxycholate. For immunoprecipitation, 50 μL of Protein G Dynabeads (Invitrogen) were pre-incubated with 2.4 μg of either CaMKIIα (Leonard et al., 1998; Leonard et al., 1999; Lu et al., 2007) or mouse IgG antibody (sc-2025, Santa Cruz Biotechnology) at RT for 10 min, washed with 0.05% TBS-tween, incubated with 1000-1500 μg of protein lysate for 30 min at RT, washed four times with 0.01% TBS-triton, and then eluted. For PSD isolation, lysates were fractionated by centrifugation and sucrose gradient, and extracted with Triton X-100, as described (Dosemeci et al., 2006). Protein samples were run on a SDS-PAGE gel at 30 mA and transferred to 0.45 um PVDF membrane for 210 min at 50 V. Blots were stained for total protein with Revert 700 Total Protein Stain Kit (LICOR). Membranes were blocked with TBS Odyssey Blocking Buffer (LICOR) and incubated overnight at 4°C with primary antibodies for GluN2B, GluN2A, GluN1, CaMKIIα (Leonard et al., 1998; Leonard et al., 1999; Lu et al., 2007), pT286 CaMKIIα (sc-12886R, Santa Cruz), synaptophysin (Sigma S5768), Cav1.2 (FP1) (Buonarati et al., 2017), or serine racemase (sc-365217, Santa Cruz). Secondary antibody (IRDye; LICOR) incubation was for 1 hour at RT and the blots scanned and analyzed using Odyssey CLx and Image Studio.

### Experimental Design and Statistical analysis

Cells for each condition were obtained from at least 3 independent hippocampal acute slices preparations of both sexes. Data analysis was done blind to the experimental condition. All statistics were calculated across cells and performed in GraphPad Prism 8.0. Student’s unpaired t-test was used for all experiments. Details on ‘n’ are included in the figure legends. All data are represented as mean ± standard error of the mean (SEM). Statistical significance was set at p < 0.05 (two-tailed t test).

## Results

### Activity-induced growth of dendritic spines is impaired in SRKO mice

NMDAR-dependent long-term potentiation (LTP) of synaptic strength and associated spine growth, mediated by simultaneous binding of glutamate and co-agonist, D-serine or glycine, is an important cellular process for memory formation and maintenance of spine density. Based on the requirement for robust calcium influx during NMDAR-dependent spine growth and stabilization, we hypothesized that reduced bioavailability of D-serine observed in individuals with schizophrenia (Bendikov et al., 2007; Hashimoto et al., 2003a) would inhibit LTP-associated spine growth. Furthermore, we predicted that the reduction in D-serine levels would increase ion flux-independent NMDAR signaling, resulting in further spine destabilization that could contribute to the reduced spine density associated with the disorder (Rosoklija et al., 2000; Sweet et al., 2009).

To begin testing our hypothesis, we first examined whether spine structural plasticity is altered in the serine racemase knockout (SRKO) mouse line, which lacks the enzyme required for D-serine production and serves as an important model for studying the consequences of reduced D-serine levels, which have been associated with schizophrenia (Basu et al., 2009). To visualize spines and dendrites for monitoring activity-dependent long-term spine growth, we crossed the SRKO mouse line with the GFP-M (Feng et al., 2000) mouse line to obtain D-serine deficient mice that have sparse neuronal GFP expression in the hippocampus. Using high-frequency uncaging (HFU) of glutamate to induce long-term spine growth at single dendritic spines on basal dendrites of CA1 neurons in acute hippocampal slices, we observed a complete inhibition of long-term spine growth in SRKO mice relative to WT littermates (**Fig. 1A-C;** WT: 168 ± 17%; KO: 103 ± 7%). Notably, the lack of long-term spine growth in SRKO was rescued with acute treatment of 10 μM D-serine (**Fig. 1D-F;** WT: 201 ± 23%; KO: 169 ± 16%), demonstrating that the deficit in long-term spine growth is not due to chronic alterations in synaptic function in SRKO animals or a role for serine racemase independent of D-serine synthesis. Thus, aligned with the previously reported LTP deficit in these mice (Basu et al., 2009), we conclude that activity-induced spine growth is impaired in SRKO mice.

**Figure 1.**
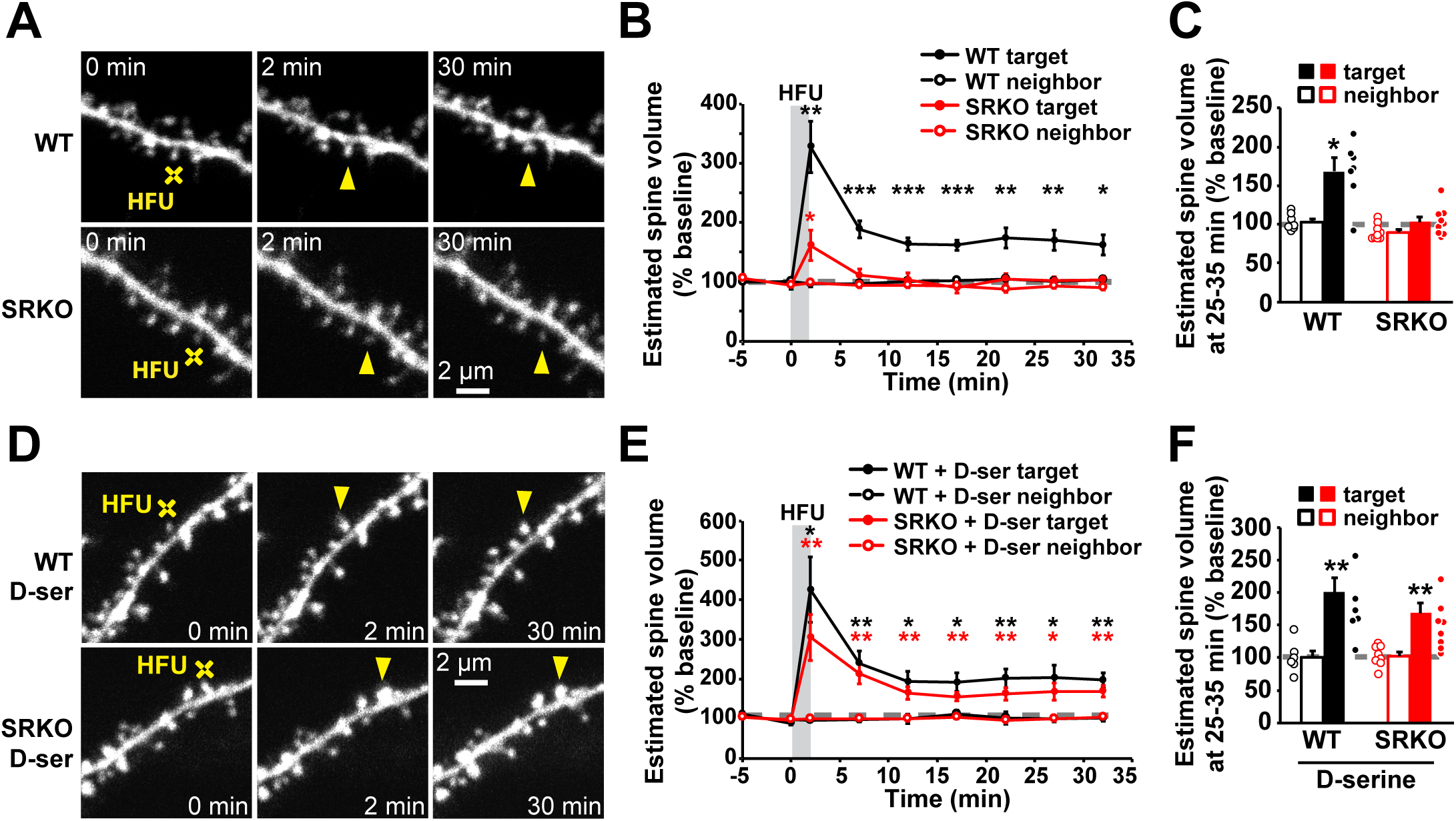
LTP-associated growth of dendritic spines is impaired in SRKO mice. **(A, D)** Images of basal dendrites of CA1 pyramidal neurons in acute hippocampal slices from WT and SRKO mice (P14-21). Individual spines (yellow arrowhead) were stimulated with high frequency glutamate uncaging (HFU, yellow crosshair) during vehicle condition or in the presence of D-serine (10 μM). (**B, C)** HFU leads to long-term spine growth in WT (black filled circles/bar; n=7 cells/7 mice; p=0.004) but not in SRKO (red filled circles/bar; n=9 cells/8 mice; p=0.63). Volume of unstimulated neighboring spines were not affected (open circles/bars). **(E, F)** Addition of D-serine rescues HFU-induced long-term spine growth (red filled circles/bar; n=8 cells/7 mice; p=0.005) to levels comparable to those in WT (black filled circles/bar; n=6 cells/5 mice; p=0.007). Data are represented as mean +/- SEM. *p<0.05; **p<0.01; ***p<0.001.

### Shift toward spine destabilization in SRKO mice

Because we observed a complete block of activity-dependent spine growth in the SRKO mice, we further wondered whether the reduced D-serine levels generated abnormal conditions, beyond just reducing ion flow through NMDARs, that disrupted downstream signaling and completely inhibited all forms of spine structural plasticity. We therefore tested whether increasing or decreasing the influx of Ca^2+^ through the NMDAR would be sufficient to drive downstream signaling and spine structural plasticity in the SRKO.

We first tested whether increasing the amount of Ca^2+^ influx would overcome the impairment of activity-dependent spine growth in the SRKO. We increased the extracellular Ca^2+^ concentration to 3 mM in order to strengthen the influx of Ca^2+^ in response to HFU stimulation, while maintaining a set amount of glutamate binding to the NMDAR. We found that HFU in 3 mM Ca^2+^ induced long-term spine growth in both WT and SRKO mice (**Fig. 2A, B;** WT: 187 ± 25%; KO: 193 ± 35%), demonstrating that the signaling mechanisms downstream of Ca^2+^ influx that support long-term spine growth are intact in SRKO mice.

**Figure 2.**
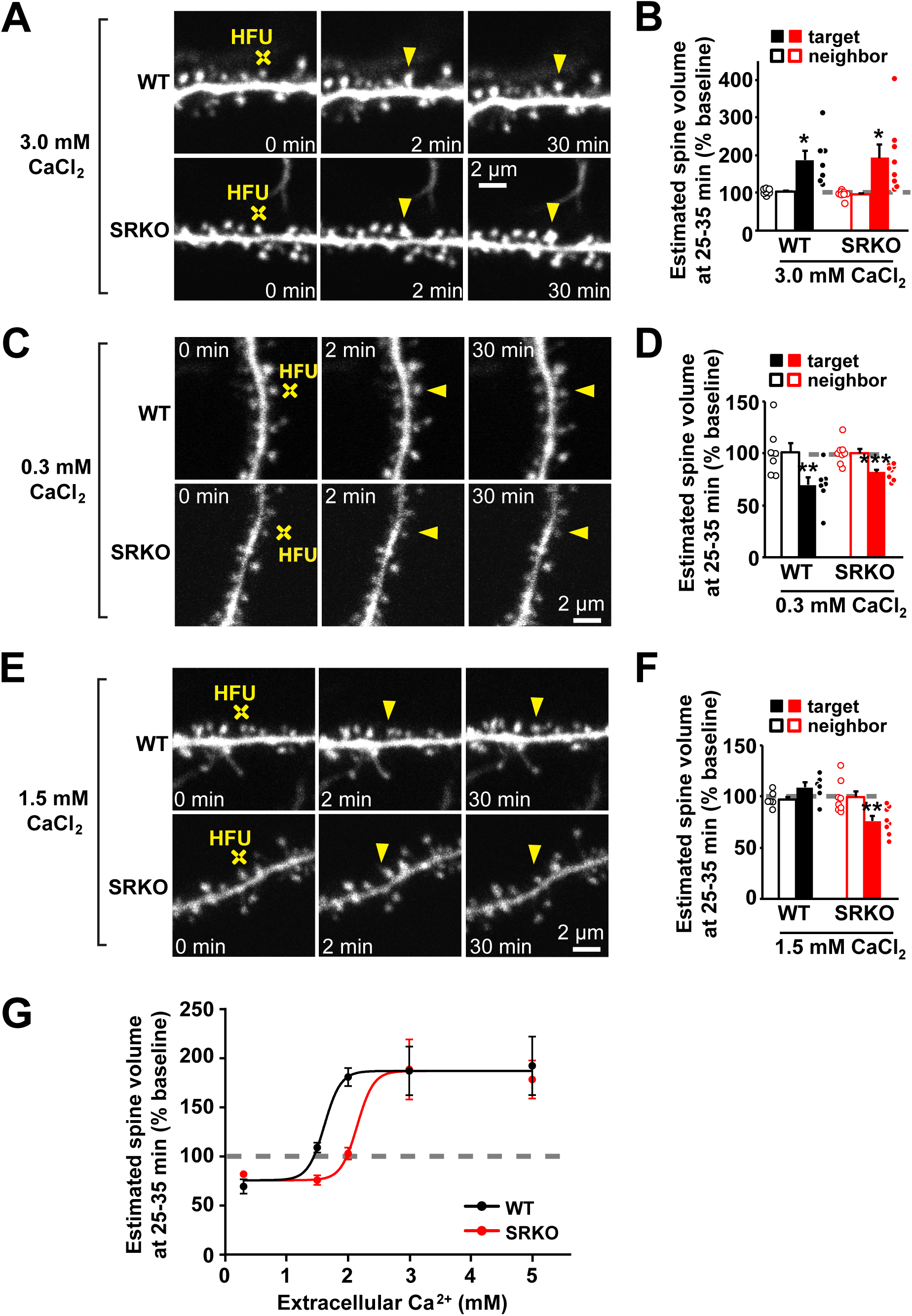
Structural plasticity is shifted to favor spine shrinkage in SRKO mice. **(A,C,E)** Images of basal dendrites of CA1 pyramidal neurons from WT and SRKO mice (P14-21). Individual spines (yellow arrowhead) were stimulated with HFU (yellow crosshair) in ACSF with different CaCl_2_ concentrations. **(B)** HFU in ASCF containing 3 mM CaCl_2_ drives long-term spine growth in both WT (black filled circles/bar; n=7 cells/5 mice; p=0.013) and SRKO (red filled circles/bar; n=9 cells/3 mice; p=0.020). **(D)** HFU in ACSF containing 0.3 mM CaCl_2_ drives long-term spine shrinkage in both WT (black filled circles/bar; n=7 cells/4 mice; p=0.0061) and SRKO (red filled circles/bar; n=8 cells/5 mice; p=0.0002). **(F)** HFU in ACSF containing 1.5 mM CaCl_2_ does not drive any long-term spine volume change in WT (black filled circles/bar; n=6 cells/6 mice; p=0.13) but drives shrinkage in SRKO (red filled circles/bar; n=8 cells/5 mice; p=0.001). **(G)** Structural plasticity curve of SRKO mice is shifted to the right, demonstrating a bias for spine shrinkage. Data are represented as mean +/- SEM. *p<0.05; **p<0.01; ***p<0.001.

We next tested whether decreasing the amount of Ca^2+^ influx would result in activity-dependent spine shrinkage in the SRKO. We decreased the extracellular Ca^2+^ concentration to 0.3 mM in order to weaken the influx of Ca^2+^ in response to HFU stimulation, while maintaining a set amount of glutamate binding to the NMDAR. We found that HFU in 0.3 mM Ca^2+^ induced long-term spine shrinkage in both WT and SRKO mice (**Fig. 2C, D;** WT: 69 ± 7%; KO: 82 ± 3%), demonstrating that the signaling mechanisms that support long-term spine shrinkage are intact in SRKO mice.

Because we found bidirectional spine structural plasticity intact in SRKO mice, we hypothesized that the disruption of activity-dependent long-term spine growth in SRKO mice that we observed in **Fig. 1** was due to a bias in spine structural plasticity toward shrinkage relative to that of WT mice. We therefore predicted that a Ca^2+^ concentration that resulted in no spine structural plasticity (at the threshold between spine shrinkage and growth) in WT mice would drive spine shrinkage in the SRKO mice. Indeed, we found that HFU in 1.5 mM Ca^2+^, which did not lead to spine structural plasticity in WT mice, resulted in spine shrinkage in SRKO mice (**Fig. 2E, F;** WT: 109 ± 5%; KO: 76 ± 5%). Plotting all of our data together, we observed that both genotypes have similar S-shaped plasticity curves, but the SRKO plasticity curve is shifted to the right (**Fig. 2G**), demonstrating that a wider range of Ca^2+^ levels elicit spine shrinkage in the SRKO, thus biasing spine structural plasticity toward shrinkage.

### Increased synaptic NMDARs and decreased CaMKII activation in SRKO

Recent studies have established that glutamate binding to the NMDAR in the absence of co-agonist binding is sufficient to drive destabilization and shrinkage of dendritic spines (Stein et al., 2015; Stein et al., 2020; Thomazeau et al., 2020). We predicted that binding of glutamate to NMDAR in the absence of co-agonist would drive ion-flux independent NMDAR signaling and contribute to spine destabilization and shrinkage in SRKO mice. Furthermore, because adult SRKO mice exhibit elevated NMDAR expression (Balu and Coyle, 2011; Mustafa et al., 2010), we wondered if NMDARs were also elevated in hippocampus in young mice and, in combination with the reduction in D-serine levels, would contribute to even further increase in non-ionotropic NMDAR signaling. Thus, we hypothesized that the combination of reduced D-serine levels and increased expression of NMDARs available for non-ionotropic NMDAR signaling led to the observed bias for spine shrinkage in SRKO.

To investigate whether SRKO mice also have increased synaptic NMDAR expression levels at dendritic spines within the hippocampus, we isolated postsynaptic density (PSD) fractions from the hippocampi of P20 SRKO mice. We found an increased levels of the obligatory NMDAR subunit GluN1 in SRKO animals relative to WT littermates, suggesting greater number of synaptic NMDARs (**Fig. 3A, B;** KO: 141 ± 11%). Interestingly, we also observed increased synaptic enrichment (**Fig. 3A, B;** KO: 237 ± 15%) and total expression (**Fig. 3C, D;** KO: 122 ± 3%) of GluN2B in SRKO relative to WT. Despite the greater number of NMDARs, we expected disrupted calcium-dependent NMDAR signaling due to reduced ion flux through the NMDAR. Indeed, when we probed CaMKII-GluN2B interaction by immunoprecipitation or phosphorylation of CaMKII at the T286 autophosphorylation site, both induced by strong calcium influx and integral for LTP (Halt et al., 2012; Lee et al., 2009), we found that basal levels of CaMKII-GluN2B interaction and pT286 are both decreased in SRKO compared to WT (**Fig. 3E, F;** KO CaMKII-GluN2B: 68 ± 6%; KO pT286: 89 ± 1%), despite no change in CaMKII expression or enrichment levels (**Fig. 3A, B;** KO CaMKII expression: 112 ± 4%; KO synaptic enrichment: 114 ± 15%). In sum, our results confirm increased NMDAR content and decreased calcium-dependent NMDAR downstream signaling in hippocampal synapses of SRKO mice.

**Figure 3.**
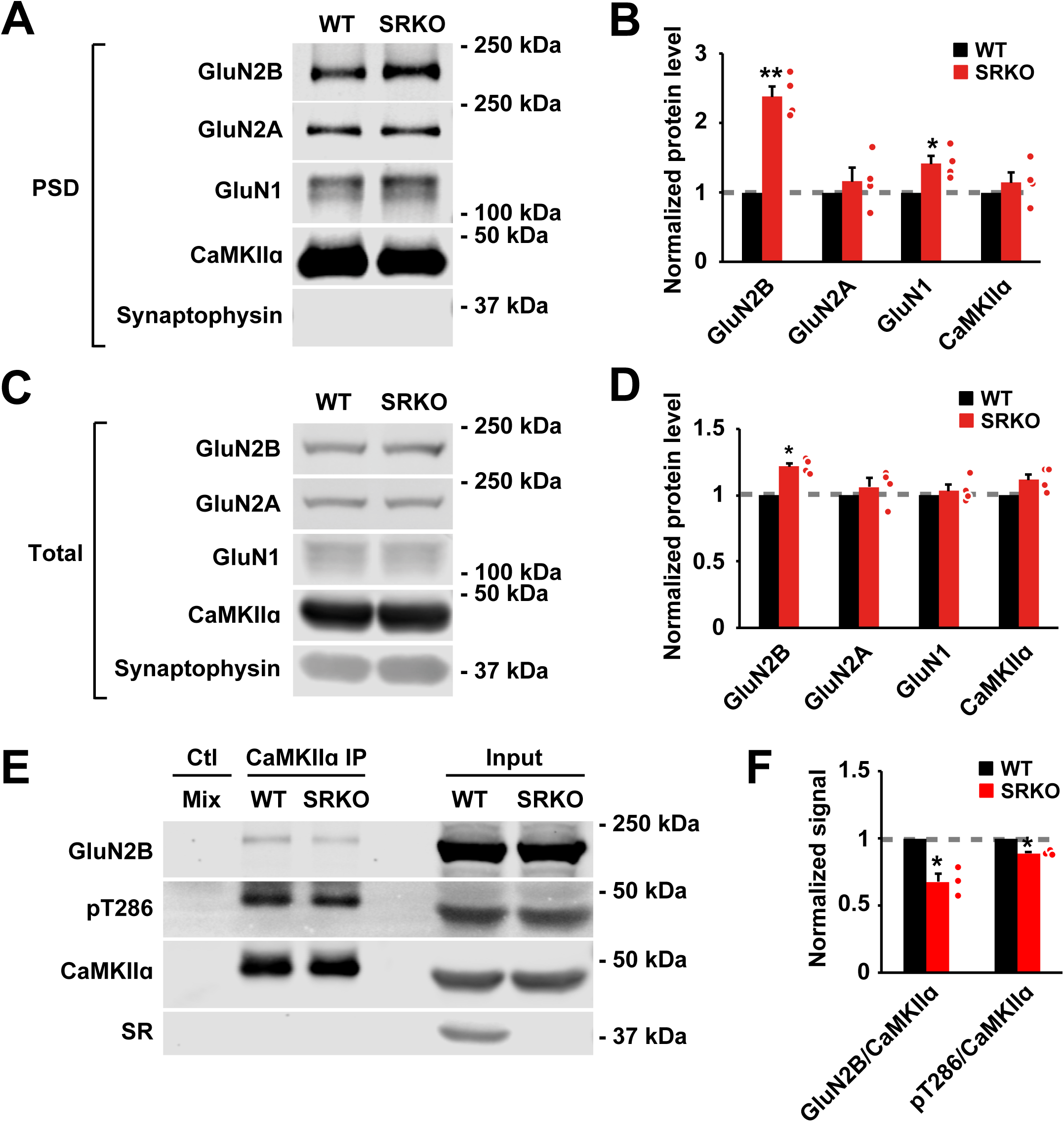
Increased synaptic NMDARs and decreased CaMKII activation in SRKO mice. **(A, B)** PSD signals from P20 SRKO hippocampi (n=12 mice/4 preps) show increased levels of synaptic GluN2B (p=0.003) and GluN1 (p=0.033) relative to WT. No change in levels of GluN2A (p=0.49) or CaMKII (p=0.41) was observed. **(C, D)** Total homogenate (n=4 mice/ 4 preps) signal shows increased levels of GluN2B (p=0.004) relative to WT. No change in levels of GluN2A (p=0.41), GluN1 (p=0.50), or CaMKII (p=0.074) were observed. **(E, F)** Immunoprecipitation of CaMKII from P20 SRKO hippocampi (n=3 mice/3 preps) shows decreased CaMKII-GluN2B interaction (p=0.034) and decreased pT286 levels of CaMKII (p=0.010) relative to WT. Data are represented as mean +/- SEM. *p<0.05; **p<0.01; ***p<0.001.

### Enhanced non-ionotropic NMDAR signaling in SRKO mice

Our observations of increased number of synaptic NMDARs in mice with reduced D-serine levels support that altered structural plasticity in SRKO could be due to increased non-ionotropic NMDAR signaling driven by glutamate binding to the increased number of NMDARs in the absence of co-agonist binding. Because there is no direct means of measuring the relative amount of non-ionotropic signaling, we first set out to probe the relative contribution of non-ionotropic NMDAR signaling to spine structural plasticity between the SRKO and WT using an assay that assesses non-ionotropic NMDAR signaling levels by monitoring spine growth that is driven with calcium influx through voltage-gated calcium channels (VGCCs) (Stein et al., 2021) (**Fig. 4A**). We hypothesized that, under conditions of robust calcium influx through VGCCs, we would observe enhanced spine growth in the SRKO due to the enhanced synaptic NMDARs driving more non-ionotropic NMDAR signaling.

**Figure 4.**
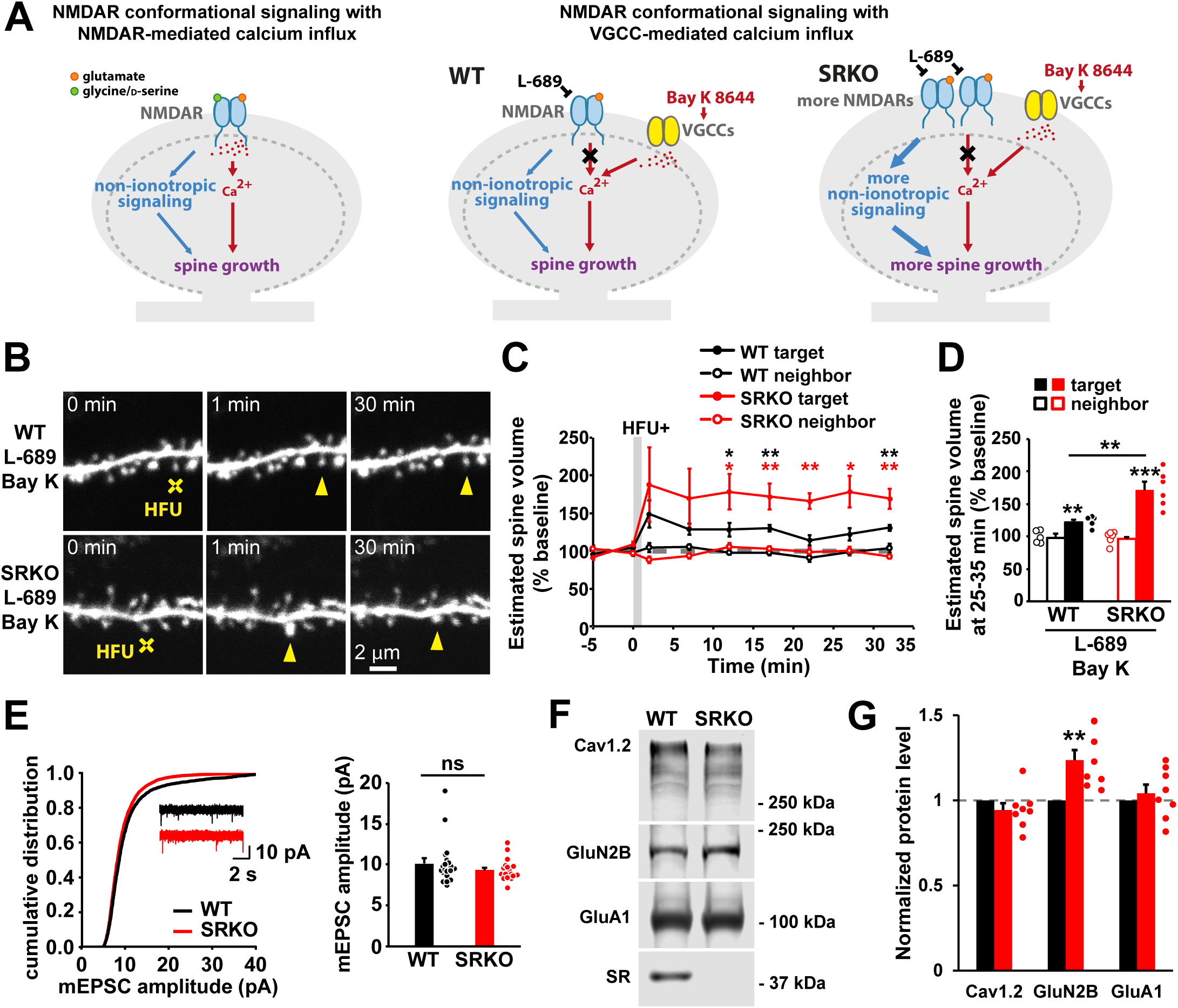
Enhanced non-ionotropic NMDAR signaling in SRKO mice. **(A)** Left: Schematic of the signaling requirements for long-term spine growth, which include non-ionotropic NMDAR signaling and NMDAR-mediated Ca^2+^ influx. Right: Experimental design for assessing levels of non-ionotropic NMDAR signaling in WT versus SRKO mice and predicted outcomes. **(B)** Images of dendrites before and after HFU+ stimulation (yellow crosshair) at individual spines (yellow arrowhead) in the presence of L689 (10 μM), an NMDAR co-agonist site inhibitor used to block Ca^2+^ influx through NMDARs, and Bay K (10 μM) to promote Ca^2+^ influx through VGCCs. **(C, D)** Although HFU+ stimulation drives long-term spine growth in WT (black filled circles/bar; n=5 cells/5 mice; p=0.005) and SRKO (red filled circles/bar; n=6 cells/4 mice; p=0.001), the magnitude of long-term spine growth is larger in SRKO than WT (p=0.006). **(E)** Amplitude of mEPSCs of P15-19 CA1 pyramidal neurons is unaltered in SRKO (WT: black line/bar; n=20 cells/3 mice; SRKO red line/bar: n=17 cells/3 mice; p=0.26). **(F, G)** Synaptosomal signal from P20 SRKO hippocampi shows no change in GluA1 (p=0.22) or Cav1.2 (p=0.10) from WT levels, despite increased levels of GluN2B relative to WT (n=7 preps/21 mice, p = 0.003). Data are represented as mean +/- SEM. *p<0.05; **p<0.01; ***p<0.001

To isolate non-ionotropic NMDAR signaling in SRKO mice, we used the NMDAR co-agonist site inhibitor L-689,560 (L-689) to mimic the absence of co-agonist and thus block NMDAR-mediated Ca^2+^ influx following glutamate binding. We combined this with the L-type Ca^2+^ channel agonist Bay K 8644 (Bay K) to promote Ca^2+^ influx through VGCCs. A modified HFU paradigm (HFU+) was used to induce a weak, non-saturated long-term increase in spine size in WT mice (**Fig. 4B-D**). Remarkably, the same HFU+ stimulation drove a robust increase in spine growth in SRKO mice (**Fig. 4B-D;** WT: 122 ± 4%; KO: 171 ± 11%). The enhancement in spine growth cannot be attributed to increased AMPAR function in SRKO mice, as the amplitude of miniature excitatory postsynaptic currents (mEPSCs) was not different between WT and SRKO (**Fig. 4E;** WT: 10 ± 0.6 pA; KO: 9.3 ± 0.3 pA), nor can it be attributed to increased expression of synaptic voltage-gated calcium channels or AMPARs, as synaptosomal preparations showed no change in Cav1.2 or GluA1 in SRKO compared to WT, despite the increased GluN2B levels (**Fig. 4F, G**; KO Cav1.2: 94 ± 4%; KO GluA1: 104 ± 5%; KO GluN2B: 124 ± 6%). We therefore interpret the greater amount of spine growth in SRKO to be indicative of a higher level of non-ionotropic NMDAR signaling.

To confirm using a different approach that increased non-ionotropic NMDAR signaling in SRKO causes a shift of spine plasticity toward shrinkage, we next used L689 to block the co-agonist binding site during a weak glutamate-uncaging stimulation. We reasoned that if SRKO mice have more synaptic NMDARs and thus undergo more non-ionotropic NMDAR signaling relative to WT, a weak glutamate uncaging stimulation that fails to induce spine shrinkage in WT should be sufficient to drive shrinkage in SRKO. Indeed, while our weaker HFU stimulus (reduced pulse width of 1 ms) in the presence of L689 and NBQX failed to induce shrinkage of dendritic spines in WT animals (**Fig. 5A-C**; WT 1 ms: 99 ± 2%), despite robust shrinkage observed with the control HFU 2 ms stimulus (**Fig. 5A-C**; WT 2 ms: 66 ± 5%), it was sufficient to drive shrinkage of dendritic spines in SRKO mice (**Fig. 5A-C**; SRKO 1 ms: 72 ± 4%). These combined results support our model (**Fig. 5D**) in which conditions of reduced D-serine levels in SRKO lead to increased synaptic NMDARs, which drive enhanced non-ionotropic NMDAR signaling, and thus create a bias toward spine shrinkage and loss.

**Figure 5.**
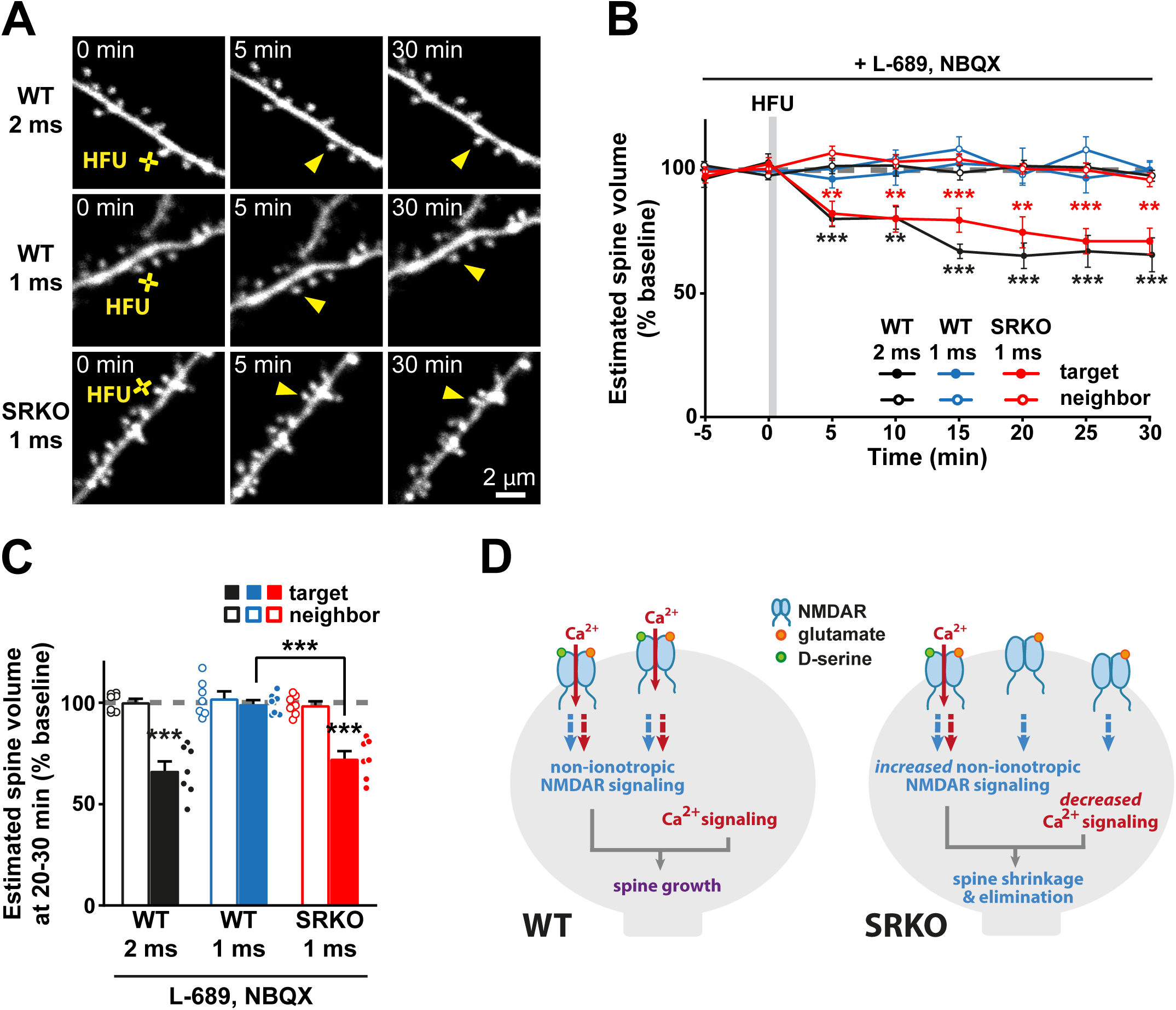
Shift of plasticity toward spine shrinkage in SRKO mice. **(A)** Images of basal dendrites of CA1 pyramidal neurons from P14-21 WT and SRKO mice. Individual spines (yellow arrowhead) were stimulated with HFU (yellow crosshair) in the presence of L-689 (10 μM) and NBQX (50 μM) for 1 or 2 ms. **(B, C)** HFU in L-689 with 2 ms pulse duration (black filled circles/bar; n=7 cells/5 mice; p=0.0006) leads to shrinkage of WT target spines, but not HFU in L-689 with 1 ms pulse duration (blue filled circles/bar; n=7 cells/5 mice; p=0.6). In contrast, HFU in L-689 with 1 ms pulse duration (red filled circles/bar; n=7 cells/6 mice; p=0.0006) is sufficient to drive spine shrinkage in SRKO mice. Neighboring spines were unchanged in all conditions (open circles/bars). Data are represented as mean +/- SEM. *p<0.05; **p<0.01; ***p<0.001 **(D)** Proposed model. Binding of glutamate and D-serine leads to both non-ionotropic NMDAR signaling and calcium influx required for activity-induced long-term spine growth. In contrast, increased NMDAR levels and reduced levels of D-serine in the SRKO mouse model for studying schizophrenia results in glutamate binding alone to more NMDARs in the absence of D-serine that results in strong non-ionotropic NMDAR activation with small calcium influx, promotes spine shrinkage and destabilization.

## Discussion

### Lowered D-serine levels create a bias towards spine shrinkage

It has been widely reported that schizophrenia is associated with a reduction in dendritic spine density that is thought to contribute to cognitive deficits. As changes in spine density and gray matter volume mirror each other (Bennett, 2011), and longitudinal MRI studies of high risk individuals report normal increase in gray matter during childhood that then declines in adolescence (Job et al., 2005; Pantelis et al., 2003; Thompson et al., 2001), a time when spine pruning increases (Penzes et al., 2011), it has been suggested that excessive spine elimination, rather than a deficit in new spine outgrowth, is the potential cause of decreased spine density in schizophrenia (Glausier and Lewis, 2013).

Here, we show a disruption of long-term spine growth and a bias toward activity-induced spine shrinkage in the SRKO mouse model for studying schizophrenia, which is reported to have decreased spine density at older ages (Balu et al., 2013). Although we observed a shift in the plasticity curve for the SRKO, we were able to induce both spine shrinkage and growth, and it has been demonstrated that spine density can be rescued in these mice with D-serine treatment (Balu and Coyle, 2014). Thus, our studies support that serine racemase, the enzyme that produces D-serine and observed to interact with various synaptic structural proteins such as PSD95 (Lin et al., 2016; Ma et al., 2014), is itself not required for structural plasticity of spines. Notably, D-serine has been shown to promote spine stability (Lin et al., 2016), likely through increased incidence of simultaneous glutamate and co-agonist binding to the NMDAR for Ca^2+^ influx required for stabilization (Hill and Zito, 2013). As glutamate binding alone to the NMDAR drives spine shrinkage in an ion-flux independent manner (Stein et al., 2015), we propose that enhancement of non-ionotropic NMDAR signaling due to the decreased levels of D-serine in SRKO mice biases toward spine shrinkage and may also contribute to spine loss associated with schizophrenia.

### Enhanced levels and altered composition of synaptic NMDARs in SRKO mice

Our observation of increased synaptic enrichment of NMDARs relative to WT within the hippocampus of young mice is consistent with previous studies on SRKO mice showing increased expression of GluN1 (Balu and Coyle, 2011; Mustafa et al., 2010) and GluN2B (Basu et al., 2009; Wong et al., 2020). These changes in the overall number and composition of NMDARs may be a direct consequence of reduced D-serine levels, as prior studies have demonstrated the role of co-agonist binding in priming of the NMDAR for endocytosis, with D-serine specifically acting on GluN2B subunits (Ferreira et al., 2017; Nong et al., 2003). In addition, we found that this increase in the number of NMDARs led to an increase in the magnitude of non-ionotropic signaling in SRKO, which would be expected to drive enhanced spine destabilization and shrinkage (Stein et al., 2015; Stein et al., 2021). Notably, despite the enhancement of NMDAR levels at the synapse, we observed decreased CaMKII-GluN2B interaction and decreased autophosphorylation of CaMKII in SRKO mice, which we attribute to the lack of strong Ca^2+^ influx required for increasing both CaMKII activity and interaction to GluN2B (Goodell et al., 2017). This altered downstream CaMKII signaling likely contributes to NMDAR hypofunction in schizophrenia (Banerjee et al., 2015).

### Enhanced non-ionotropic NMDAR signaling in SRKO mice

We made several observations that support our hypothesis that there is increased ion flux-independent NMDAR signaling driving spine destabilization in the SRKO mice. First, we observed that stimulation protocols which normally induce spine growth in WT mice instead drive spine shrinkage in SRKO. Second, we found increased NMDARs at hippocampal synapses in the SRKO mice, which, in combination with the reduced D-serine levels, biases NMDARs toward ion flux-independent signaling. Third, we showed increased ion flux-independent NMDAR signaling in SRKO animals. Finally, we observed that weak activation of non-ionotropic NMDAR signaling that fails to induce spine shrinkage in WT can drive spine shrinkage in SRKO mice, further supporting enhanced non-ionotropic NMDAR signaling in SRKO mice. This enhanced non-ionotropic NMDAR signaling drives spine destabilization and shrinkage and would be expected to drive eventual spine elimination.

Notably, studies in which NMDAR channel blockers produce schizophrenia-like symptoms in healthy individuals and exacerbate them in individuals with schizophrenia helped give rise to the NMDAR hypofunction hypothesis (Javitt and Zukin, 1991; Krystal et al., 1994; Lahti et al., 2001; Newcomer et al., 1999). Due to high sensitivity of inhibitory GABAergic neurons to NMDAR blockers (Grunze et al., 1996; Homayoun and Moghaddam, 2007), decreased expression of interneurons (Hashimoto et al., 2008; Hashimoto et al., 2003b; Mellios et al., 2009), and GABAergic markers (Glausier and Lewis, 2017; Gonzalez-Burgos et al., 2011; Lewis et al., 2008; Lewis et al., 1999), NMDAR hypofunction caused by reduced D-serine levels in schizophrenia may lead to disinhibition of excitatory neurons and result in glutamate spillover (Gallinat et al., 2016; Kraguljac et al., 2013; Lorrain et al., 2003; van Elst et al., 2005). Indeed, SRKO mice have been observed to have decreased PV expression and altered excitatory/inhibitory balance from GABAergic dysfunction (Jami et al., 2020; Ploux et al., 2020; Steullet et al., 2017). This disinhibition should result in greater release of glutamate at dendritic spines that, when paired with reduced D-serine levels and increased levels of synaptic NMDARs, would increase the amount of non-ionotropic NMDAR activation even further to promote spine shrinkage (Stein et al., 2020) and decrease in spine density in the SRKO and schizophrenia (Balu et al., 2013; Rosoklija et al., 2000; Sweet et al., 2009).

Here we show that the SRKO mouse model for studying schizophrenia displays altered dendritic structural plasticity that biases toward spine shrinkage. We further report an increased number of synaptic NMDARs in the hippocampus of the SRKO mice and increased ion flux-independent NMDAR signaling at hippocampal spines. Taken together, our findings support a model in which NMDAR hypofunction brought on by lack of D-serine, promotes excessive non-ionotropic NMDAR signaling that drives a bias towards spine shrinkage and likely contributes to decreased spine density associated with schizophrenia.

## Author contributions

D.K.P., I.S.S., and K.Z. designed the study and wrote the initial draft of the manuscript. D.K.P., S.P., and M.A. performed imaging and biochemistry experiments and analysis. D.K.P., J.A.G., and E.V.B. designed and E.V.B. performed electrophysiology experiments and analysis. All authors edited the manuscript.

## Acknowledgements

This work was supported by the NIH (R01 NS062736, R01 MH117130 T32 GM007377, T32 MH112507, T32 GM099608), an ARCS scholar award (D.K.P.), and the Deutsche Forschungsgemeinschaft Walter Benjamin project 468470832 (M.A.). We thank Joseph Coyle for the SRKO mice; Johannes Hell for CaMKIIα, GluN1, GluN2B, GluN2A, GluA1, and Cav1.2 antibodies; Julie Culp, Jennifer Jahncke, and Lorenzo Tom for support with experiments and analysis; and Joseph Coyle, Darrick Balu, Johannes Hell, and Scott Cameron for critical input.

## Conflict of interest

The authors declare no competing interests.

## Notes

### Competing Interest Statement

The authors have declared no competing interest.

### Summary of Updates

updated title

## References

Alvarez, V.A., Ridenour, D.A., and Sabatini, B.L. (2007). Distinct structural and ionotropic roles of NMDA receptors in controlling spine and synapse stability. J Neurosci 27, 7365–7376.

Balu, D.T., and Coyle, J.T. (2011). Glutamate receptor composition of the post-synaptic density is altered in genetic mouse models of NMDA receptor hypo- and hyperfunction. Brain Res 1392, 1–7.

Balu, D.T., and Coyle, J.T. (2014). Chronic D-serine reverses arc expression and partially rescues dendritic abnormalities in a mouse model of NMDA receptor hypofunction. Neurochem Int 75, 76–78.

Balu, D.T., Li, Y., Puhl, M.D., Benneyworth, M.A., Basu, A.C., Takagi, S., Bolshakov, V.Y., and Coyle, J.T. (2013). Multiple risk pathways for schizophrenia converge in serine racemase knockout mice, a mouse model of NMDA receptor hypofunction. Proc Natl Acad Sci U S A 110, E2400–2409.

Banerjee, A., Wang, H.Y., Borgmann-Winter, K.E., MacDonald, M.L., Kaprielian, H., Stucky, A., Kvasic, J., Egbujo, C., Ray, R., Talbot, K., et al. (2015). Src kinase as a mediator of convergent molecular abnormalities leading to NMDAR hypoactivity in schizophrenia. Mol Psychiatry 20, 1091–1100.

Barnes, S.A., Sawiak, S.J., Caprioli, D., Jupp, B., Buonincontri, G., Mar, A.C., Harte, M.K., Fletcher, P.C., Robbins, T.W., Neill, J.C., et al. (2014). Impaired limbic cortico-striatal structure and sustained visual attention in a rodent model of schizophrenia. Int J Neuropsychopharmacol 18.

Basu, A.C., Tsai, G.E., Ma, C.L., Ehmsen, J.T., Mustafa, A.K., Han, L., Jiang, Z.I., Benneyworth, M.A., Froimowitz, M.P., Lange, N., et al. (2009). Targeted disruption of serine racemase affects glutamatergic neurotransmission and behavior. Mol Psychiatry 14, 719–727.

Bendikov, I., Nadri, C., Amar, S., Panizzutti, R., De Miranda, J., Wolosker, H., and Agam, G. (2007). A CSF and postmortem brain study of D-serine metabolic parameters in schizophrenia. Schizophr Res 90, 41–51.

Bennett, M.R. (2011). Schizophrenia: susceptibility genes, dendritic-spine pathology and gray matter loss. Prog Neurobiol 95, 275–300.

Bischofberger, J., Engel, D., Li, L., Geiger, J.R., and Jonas, P. (2006). Patch-clamp recording from mossy fiber terminals in hippocampal slices. Nat Protoc 1, 2075–2081.

Brigman, J.L., Wright, T., Talani, G., Prasad-Mulcare, S., Jinde, S., Seabold, G.K., Mathur, P., Davis, M.I., Bock, R., Gustin, R.M., et al. (2010). Loss of GluN2B-containing NMDA receptors in CA1 hippocampus and cortex impairs long-term depression, reduces dendritic spine density, and disrupts learning. J Neurosci 30, 4590–4600.

Buonarati, O.R., Henderson, P.B., Murphy, G.G., Horne, M.C., and Hell, J.W. (2017). Proteolytic processing of the L-type Ca (2+) channel alpha 11.2 subunit in neurons. F1000Res 6, 1166.

Coyle, J.T. (2017). Schizophrenia: Basic and Clinical. Adv Neurobiol 15, 255–280.

Dosemeci, A., Tao-Cheng, J.H., Vinade, L., and Jaffe, H. (2006). Preparation of postsynaptic density fraction from hippocampal slices and proteomic analysis. Biochem Biophys Res Commun 339, 687–694.

Feng, G., Mellor, R.H., Bernstein, M., Keller-Peck, C., Nguyen, Q.T., Wallace, M., Nerbonne, J.M., Lichtman, J.W., and Sanes, J.R. (2000). Imaging neuronal subsets in transgenic mice expressing multiple spectral variants of GFP. Neuron 28, 41–51.

Ferreira, J.S., Papouin, T., Ladepeche, L., Yao, A., Langlais, V.C., Bouchet, D., Dulong, J., Mothet, J.P., Sacchi, S., Pollegioni, L., et al. (2017). Co-agonists differentially tune GluN2B-NMDA receptor trafficking at hippocampal synapses. Elife 6.

Gallinat, J., McMahon, K., Kuhn, S., Schubert, F., and Schaefer, M. (2016). Cross-sectional Study of Glutamate in the Anterior Cingulate and Hippocampus in Schizophrenia. Schizophr Bull 42, 425–433.

Glausier, J.R., and Lewis, D.A. (2013). Dendritic spine pathology in schizophrenia. Neuroscience 251, 90–107.

Glausier, J.R., and Lewis, D.A. (2017). GABA and schizophrenia: Where we stand and where we need to go. Schizophr Res 181, 2–3.

Gonzalez-Burgos, G., Fish, K.N., and Lewis, D.A. (2011). GABA neuron alterations, cortical circuit dysfunction and cognitive deficits in schizophrenia. Neural Plast 2011, 723184.

Goodell, D.J., Zaegel, V., Coultrap, S.J., Hell, J.W., and Bayer, K.U. (2017). DAPK1 Mediates LTD by Making CaMKII/GluN2B Binding LTP Specific. Cell Rep 19, 2231–2243.

Grunze, H.C., Rainnie, D.G., Hasselmo, M.E., Barkai, E., Hearn, E.F., McCarley, R.W., and Greene, R.W. (1996). NMDA-dependent modulation of CA1 local circuit inhibition. J Neurosci 16, 2034–2043.

Halt, A.R., Dallapiazza, R.F., Zhou, Y., Stein, I.S., Qian, H., Juntti, S., Wojcik, S., Brose, N., Silva, A.J., and Hell, J.W. (2012). CaMKII binding to GluN2B is critical during memory consolidation. EMBO J 31, 1203–1216.

Hashimoto, K., Fukushima, T., Shimizu, E., Komatsu, N., Watanabe, H., Shinoda, N., Nakazato, M., Kumakiri, C., Okada, S., Hasegawa, H., et al. (2003a). Decreased serum levels of D-serine in patients with schizophrenia: evidence in support of the N-methyl-D-aspartate receptor hypofunction hypothesis of schizophrenia. Arch Gen Psychiatry 60, 572–576.

Hashimoto, T., Arion, D., Unger, T., Maldonado-Aviles, J.G., Morris, H.M., Volk, D.W., Mirnics, K., and Lewis, D.A. (2008). Alterations in GABA-related transcriptome in the dorsolateral prefrontal cortex of subjects with schizophrenia. Mol Psychiatry 13, 147–161.

Hashimoto, T., Volk, D.W., Eggan, S.M., Mirnics, K., Pierri, J.N., Sun, Z., Sampson, A.R., and Lewis, D.A. (2003b). Gene expression deficits in a subclass of GABA neurons in the prefrontal cortex of subjects with schizophrenia. J Neurosci 23, 6315–6326.

Hayashi-Takagi, A., Yagishita, S., Nakamura, M., Shirai, F., Wu, Y.I., Loshbaugh, A.L., Kuhlman, B., Hahn, K.M., and Kasai, H. (2015). Labelling and optical erasure of synaptic memory traces in the motor cortex. Nature 525, 333–338.

Hill, T.C., and Zito, K. (2013). LTP-induced long-term stabilization of individual nascent dendritic spines. J Neurosci 33, 678–686.

Homayoun, H., and Moghaddam, B. (2007). NMDA receptor hypofunction produces opposite effects on prefrontal cortex interneurons and pyramidal neurons. J Neurosci 27, 11496–11500.

Jami, S.A., Cameron, S., Wong, J.M., Daly, E.R., McAllister, A.K., and Gray, J.A. (2020). Increased excitation-inhibition balance due to a loss of GABAergic synapses in the serine racemase knockout model of NMDA receptor hypofunction. bioRxiv.

Javitt, D.C., and Zukin, S.R. (1991). Recent advances in the phencyclidine model of schizophrenia. Am J Psychiatry 148, 1301–1308.

Job, D.E., Whalley, H.C., Johnstone, E.C., and Lawrie, S.M. (2005). Grey matter changes over time in high risk subjects developing schizophrenia. Neuroimage 25, 1023–1030.

Kannangara, T.S., Eadie, B.D., Bostrom, C.A., Morch, K., Brocardo, P.S., and Christie, B.R. (2015). GluN2A-/-Mice Lack Bidirectional Synaptic Plasticity in the Dentate Gyrus and Perform Poorly on Spatial Pattern Separation Tasks. Cereb Cortex 25, 2102–2113.

Kraguljac, N.V., White, D.M., Reid, M.A., and Lahti, A.C. (2013). Increased hippocampal glutamate and volumetric deficits in unmedicated patients with schizophrenia. JAMA Psychiatry 70, 1294–1302.

Krystal, J.H., Karper, L.P., Seibyl, J.P., Freeman, G.K., Delaney, R., Bremner, J.D., Heninger, G.R., Bowers, M.B., Jr., and Charney, D.S. (1994). Subanesthetic effects of the noncompetitive NMDA antagonist, ketamine, in humans. Psychotomimetic, perceptual, cognitive, and neuroendocrine responses. Arch Gen Psychiatry 51, 199–214.

Lahti, A.C., Weiler, M.A., Tamara Michaelidis, B.A., Parwani, A., and Tamminga, C.A. (2001). Effects of ketamine in normal and schizophrenic volunteers. Neuropsychopharmacology 25, 455–467.

Latysheva, N.V., and Raevskii, K.S. (2003). Behavioral analysis of the consequences of chronic blockade of NMDA-type glutamate receptors in the early postnatal period in rats. Neurosci Behav Physiol 33, 123–131.

Lee, S.J., Escobedo-Lozoya, Y., Szatmari, E.M., and Yasuda, R. (2009). Activation of CaMKII in single dendritic spines during long-term potentiation. Nature 458, 299–304.

Leonard, A.S., Davare, M.A., Horne, M.C., Garner, C.C., and Hell, J.W. (1998). SAP97 is associated with the alpha-amino-3-hydroxy-5-methylisoxazole-4-propionic acid receptor GluR1 subunit. J Biol Chem 273, 19518–19524.

Leonard, A.S., Lim, I.A., Hemsworth, D.E., Horne, M.C., and Hell, J.W. (1999). Calcium/calmodulin-dependent protein kinase II is associated with the N-methyl-D-aspartate receptor. Proc Natl Acad Sci U S A 96, 3239–3244.

Lewis, D.A., Hashimoto, T., and Morris, H.M. (2008). Cell and receptor type-specific alterations in markers of GABA neurotransmission in the prefrontal cortex of subjects with schizophrenia. Neurotox Res 14, 237–248.

Lewis, D.A., Pierri, J.N., Volk, D.W., Melchitzky, D.S., and Woo, T.U. (1999). Altered GABA neurotransmission and prefrontal cortical dysfunction in schizophrenia. Biol Psychiatry 46, 616–626.

Lin, H., Jacobi, A.A., Anderson, S.A., and Lynch, D.R. (2016). D-Serine and Serine Racemase Are Associated with PSD-95 and Glutamatergic Synapse Stability. Front Cell Neurosci 10, 34.

Lorrain, D.S., Baccei, C.S., Bristow, L.J., Anderson, J.J., and Varney, M.A. (2003). Effects of ketamine and N-methyl-D-aspartate on glutamate and dopamine release in the rat prefrontal cortex: modulation by a group II selective metabotropic glutamate receptor agonist LY379268. Neuroscience 117, 697–706.

Lu, Y., Allen, M., Halt, A.R., Weisenhaus, M., Dallapiazza, R.F., Hall, D.D., Usachev, Y.M., McKnight, G.S., and Hell, J.W. (2007). Age-dependent requirement of AKAP150-anchored PKA and GluR2-lacking AMPA receptors in LTP. EMBO J 26, 4879–4890.

Ma, T.M., Paul, B.D., Fu, C., Hu, S., Zhu, H., Blackshaw, S., Wolosker, H., and Snyder, S.H. (2014). Serine racemase regulated by binding to stargazin and PSD-95: potential N-methyl-D-aspartate-alpha-amino-3-hydroxy-5-methyl-4-isoxazolepropionic acid (NMDA-AMPA) glutamate neurotransmission cross-talk. J Biol Chem 289, 29631–29641.

Matsuzaki, M., Honkura, N., Ellis-Davies, G.C., and Kasai, H. (2004). Structural basis of long-term potentiation in single dendritic spines. Nature 429, 761–766.

Mellios, N., Huang, H.S., Baker, S.P., Galdzicka, M., Ginns, E., and Akbarian, S. (2009). Molecular determinants of dysregulated GABAergic gene expression in the prefrontal cortex of subjects with schizophrenia. Biol Psychiatry 65, 1006–1014.

Mustafa, A.K., Ahmad, A.S., Zeynalov, E., Gazi, S.K., Sikka, G., Ehmsen, J.T., Barrow, R.K., Coyle, J.T., Snyder, S.H., and Dore, S. (2010). Serine racemase deletion protects against cerebral ischemia and excitotoxicity. J Neurosci 30, 1413–1416.

Nabavi, S., Kessels, H.W., Alfonso, S., Aow, J., Fox, R., and Malinow, R. (2013). Metabotropic NMDA receptor function is required for NMDA receptor-dependent long-term depression. Proc Natl Acad Sci U S A 110, 4027–4032.

Newcomer, J.W., Farber, N.B., Jevtovic-Todorovic, V., Selke, G., Melson, A.K., Hershey, T., Craft, S., and Olney, J.W. (1999). Ketamine-induced NMDA receptor hypofunction as a model of memory impairment and psychosis. Neuropsychopharmacology 20, 106–118.

Nong, Y., Huang, Y.Q., Ju, W., Kalia, L.V., Ahmadian, G., Wang, Y.T., and Salter, M.W. (2003). Glycine binding primes NMDA receptor internalization. Nature 422, 302–307.

Oh, W.C., Hill, T.C., and Zito, K. (2013). Synapse-specific and size-dependent mechanisms of spine structural plasticity accompanying synaptic weakening. Proc Natl Acad Sci U S A 110, E305–312.

Pantelis, C., Velakoulis, D., McGorry, P.D., Wood, S.J., Suckling, J., Phillips, L.J., Yung, A.R., Bullmore, E.T., Brewer, W., Soulsby, B., et al. (2003). Neuroanatomical abnormalities before and after onset of psychosis: a cross-sectional and longitudinal MRI comparison. Lancet 361, 281–288.

Penzes, P., Cahill, M.E., Jones, K.A., VanLeeuwen, J.E., and Woolfrey, K.M. (2011). Dendritic spine pathology in neuropsychiatric disorders. Nat Neurosci 14, 285–293.

Plitman, E., Iwata, Y., Caravaggio, F., Nakajima, S., Chung, J.K., Gerretsen, P., Kim, J., Takeuchi, H., Chakravarty, M.M., Remington, G., et al. (2017). Kynurenic Acid in Schizophrenia: A Systematic Review and Meta-analysis. Schizophr Bull 43, 764–777.

Ploux, E., Bouet, V., Radzishevsky, I., Wolosker, H., Freret, T., and Billard, J.M. (2020). Serine Racemase Deletion Affects the Excitatory/Inhibitory Balance of the Hippocampal CA1 Network. Int J Mol Sci 21.

Rosoklija, G., Toomayan, G., Ellis, S.P., Keilp, J., Mann, J.J., Latov, N., Hays, A.P., and Dwork, A.J. (2000). Structural abnormalities of subicular dendrites in subjects with schizophrenia and mood disorders: preliminary findings. Arch Gen Psychiatry 57, 349–356.

Schobel, S.A., Chaudhury, N.H., Khan, U.A., Paniagua, B., Styner, M.A., Asllani, I., Inbar, B.P., Corcoran, C.M., Lieberman, J.A., Moore, H., et al. (2013). Imaging patients with psychosis and a mouse model establishes a spreading pattern of hippocampal dysfunction and implicates glutamate as a driver. Neuron 78, 81–93.

Stein, I.S., Gray, J.A., and Zito, K. (2015). Non-Ionotropic NMDA Receptor Signaling Drives Activity-Induced Dendritic Spine Shrinkage. J Neurosci 35, 12303–12308.

Stein, I.S., Park, D.K., Claiborne, N., and Zito, K. (2021). Non-ionotropic NMDA receptor signaling gates bidirectional structural plasticity of dendritic spines. Cell Rep 34, 108664.

Stein, I.S., Park, D.K., Flores, J.C., Jahncke, J.N., and Zito, K. (2020). Molecular Mechanisms of Non-ionotropic NMDA Receptor Signaling in Dendritic Spine Shrinkage. J Neurosci 40, 3741–3750.

Steullet, P., Cabungcal, J.H., Coyle, J., Didriksen, M., Gill, K., Grace, A.A., Hensch, T.K., LaMantia, A.S., Lindemann, L., Maynard, T.M., et al. (2017). Oxidative stress-driven parvalbumin interneuron impairment as a common mechanism in models of schizophrenia. Mol Psychiatry 22, 936–943.

Sweet, R.A., Henteleff, R.A., Zhang, W., Sampson, A.R., and Lewis, D.A. (2009). Reduced dendritic spine density in auditory cortex of subjects with schizophrenia. Neuropsychopharmacology 34, 374–389.

Thomazeau, A., Bosch, M., Essayan-Perez, S., Barnes, S.A., De Jesus-Cortes, H., and Bear, M.F. (2020). Dissociation of functional and structural plasticity of dendritic spines during NMDAR and mGluR-dependent long-term synaptic depression in wild-type and fragile X model mice. Mol Psychiatry.

Thompson, P.M., Vidal, C., Giedd, J.N., Gochman, P., Blumenthal, J., Nicolson, R., Toga, A.W., and Rapoport, J.L. (2001). Mapping adolescent brain change reveals dynamic wave of accelerated gray matter loss in very early-onset schizophrenia. Proc Natl Acad Sci U S A 98, 11650–11655.

Ultanir, S.K., Kim, J.E., Hall, B.J., Deerinck, T., Ellisman, M., and Ghosh, A. (2007). Regulation of spine morphology and spine density by NMDA receptor signaling in vivo. Proc Natl Acad Sci U S A 104, 19553–19558.

Valbuena, S., and Lerma, J. (2016). Non-canonical Signaling, the Hidden Life of Ligand-Gated Ion Channels. Neuron 92, 316–329.

van Elst, L.T., Valerius, G., Buchert, M., Thiel, T., Rusch, N., Bubl, E., Hennig, J., Ebert, D., and Olbrich, H.M. (2005). Increased prefrontal and hippocampal glutamate concentration in schizophrenia: evidence from a magnetic resonance spectroscopy study. Biol Psychiatry 58, 724–730.

Wong, J.M., Folorunso, O.O., Barragan, E.V., Berciu, C., Harvey, T.L., Coyle, J.T., Balu, D.T., and Gray, J.A. (2020). Postsynaptic Serine Racemase Regulates NMDA Receptor Function. J Neurosci 40, 9564–9575.

Woods, G.F., Oh, W.C., Boudewyn, L.C., Mikula, S.K., and Zito, K. (2011). Loss of PSD-95 enrichment is not a prerequisite for spine retraction. J Neurosci 31, 12129–12138.

Wu, H., Wang, X., Gao, Y., Lin, F., Song, T., Zou, Y., Xu, L., and Lei, H. (2016). NMDA receptor antagonism by repetitive MK801 administration induces schizophrenia-like structural changes in the rat brain as revealed by voxel-based morphometry and diffusion tensor imaging. Neuroscience 322, 221–233.

Zhou, Q., Homma, K.J., and Poo, M.M. (2004). Shrinkage of dendritic spines associated with long-term depression of hippocampal synapses. Neuron 44, 749–757.

